# Development of a LAG-3 Immunohistochemistry Assay for Melanoma

**DOI:** 10.1101/2022.02.25.481964

**Authors:** Lori Johnson, Bryan McCune, Darren Locke, Cyrus Hedvat, John B. Wojcik, Caitlin Schroyer, Jim Yan, Krystal Johnson, Angela Sanders-Cliette, Sujana Samala, Lloye M. Dillon, Steven Anderson, Jeffrey Shuster

## Abstract

**Aims:** A robust immunohistochemistry (IHC) assay was developed to detect lymphocyte-activation gene 3 (LAG-3) expression by immune cells (ICs) in tumor tissues. LAG-3 is an immuno-oncology target with demonstrable clinical benefit, and there is a need for a standardized, well-characterized assay to measure its expression. This study aims to describe LAG-3 scoring criteria and present the specificity, sensitivity, analytical precision, and reproducibility of this assay.

**Methods:** The specificity of the assay was investigated by antigen competition and with *LAG3* knockout cell lines. A melanin pigment removal procedure was implemented to prevent melanin interference in IHC interpretation. Formalin-fixed, paraffin-embedded (FFPE) human melanoma samples with a range of LAG-3 expression levels were used to assess the sensitivity and analytical precision of the assay with a ≥1% cutoff to determine LAG-3–positivity. Interobserver and intraobserver reproducibility were evaluated with 60 samples in intralaboratory studies and 70 samples in interlaboratory studies.

**Results:** The LAG-3 IHC method demonstrated performance suitable for analysis of LAG-3 IC expression in clinical melanoma samples. The pretreatment step effectively removed melanin pigment that could interfere with interpretation. LAG-3 antigen competition and analysis of *LAG3* knockout cell lines indicated that the 17B4 antibody clone binds specifically to LAG-3. The intrarun repeatability, interday, interinstrument, interoperator, and interreagent lot reproducibility demonstrated a high scoring concordance (>95%). The interobserver and intraobserver reproducibility and overall interlaboratory and intralaboratory reproducibility also showed high scoring concordance (>90%).

**Conclusions:** We have demonstrated that the assay reliably assesses LAG-3 expression in FFPE human melanoma samples by IHC.

**Key messages:** *What is already known on this topic:* Lymphocyte-activation gene 3 (LAG-3) is an immune checkpoint receptor expressed on immune cells that limits T-cell activity and is being actively explored as a target for immunotherapy.

*What this study adds:* An immunohistochemistry assay was developed to detect the LAG-3 protein in formalin-fixed paraffin-embedded human tumor tissue specimens. This study describes scoring criteria and shows the specificity, sensitivity, analytical precision, and reproducibility of this assay as an aid to determine LAG-3 expression in melanoma patients using a ≥1% expression on immune cells threshold.

*How this study might affect research, practice or policy:* The study describes a key immuno-oncology checkpoint immunohistochemistry assay that is robust and suitable for clinical trials. The assay was used in RELATIVITY-047 (NCT03470922), a phase 2/3 clinical trial that compared combined nivolumab and relatlimab treatment with nivolumab monotherapy, to stratify patients based on the percentage of LAG-3–positive immune cells within the tumor region. This assay is also being used in several ongoing clinical trials evaluating clinical response to relatlimab.

## INTRODUCTION

Immune checkpoint inhibitor–based therapies have greatly improved clinical outcomes across multiple disease settings,[1, 2] including advanced melanoma,[3–5] non-small cell lung cancer,[6, 7] squamous cell carcinoma of the head and neck,[8, 9] and urothelial carcinoma,[10, 11] among others. However, given the multiple mechanisms of immune evasion utilized by cancer cells, inhibition of a single immune checkpoint, such as programmed death-1 (PD-1), may not be sufficient to overcome immune suppression.[12, 13] Novel immuno-oncology (I-O) combinations, including dual checkpoint inhibition, may be necessary to enhance efficacy and to improve the durability of patient responses.

Lymphocyte-activation gene 3 (LAG-3, CD223) is a cell-surface molecule expressed on activated CD4+ and CD8+ T cells, as well as other immune cells (ICs) including regulatory T cells, natural killer cells, B cells, macrophages, and dendritic cells, and is under investigation as an I-O therapy target.[13–17] The interaction of LAG-3 with its ligands, the major histocompatibility complex II (MHCII), and fibrinogen-like protein 1 (FGL-1), recently discovered as a LAG-3 ligand, initiates an inhibitory signal.[13, 18, 19] This signal can impair T-cell function, activation, and proliferation, decrease production of and response to proinflammatory cytokines, and decrease the development of memory T cells.

Preclinical data indicate that simultaneous activation of the LAG-3 and PD-1 pathways in tumor-infiltrating lymphocytes results in greater T-cell exhaustion than either pathway alone, and dual inhibition of these pathways may improve T-cell function and increase immune response.[20] Furthermore, combined therapy with anti–LAG-3 and anti–PD-1 agents in fibrosarcoma and colorectal adenocarcinoma mouse models resulted in synergistic antitumor activity.[16] The clinical efficacy of combining relatlimab, an anti– LAG-3 antibody, with nivolumab, an anti–PD-1 agent, was previously demonstrated in patients with previously untreated metastatic or unresectable melanoma by the phase 2/3 RELATIVITY-047 clinical trial (NCT03470922).[21] RELATIVITY-047 demonstrated superior progression-free survival (PFS) for relatlimab combined with nivolumab versus nivolumab monotherapy, regardless of LAG-3 expression.[21]

A robust immunohistochemistry (IHC) assay was developed to detect LAG-3 expression by ICs. The assay was used to stratify patients enrolled in RELATIVITY-047, based on the percentage of LAG-3–positive ICs with a morphological resemblance to lymphocytes relative to all nucleated cells within the tumor region (tumor cells [TCs], intratumoral stroma, and peritumoral stroma [the band of stromal elements directly contiguous with the outer tumor margin]) in samples containing ≥100 viable TCs. This assay is also being used in several ongoing clinical trials evaluating relatlimab. This study presents the specificity, sensitivity, analytical precision, and reproducibility of this assay as an aid to determine LAG-3 expression in melanoma patients using a ≥1% IC expression threshold.

## MATERIALS AND METHODS

### Principles of the LAG-3 IHC assay

The LAG-3 IHC assay was developed using a mouse monoclonal antibody clone 17B4 that was made to a synthetic peptide corresponding to the 30–amino acid extra-loop of the first immunoglobulin domain of LAG-3, GPPAAAPGHPLAPGPHPAAPSSWGPRPRRY.[22] The assay was performed on formalin-fixed paraffin-embedded (FFPE) tissue sections mounted on glass slides and included pretreatment to remove endogenous melanin that could interfere with interpretation of LAG-3 staining. Following pretreatment, slides were stained and processed using the 17B4 primary antibody on a Leica BOND-III autostainer (Leica Biosystems, Buffalo Grove, IL).

### Materials

#### Tissue specimens

FFPE melanoma specimens and control tonsil tissues were obtained from commercial vendors (Boca Biolistics, Pompano Beach, FL; BioIVT, Westbury, NY; and Avaden Biosciences, Seattle, WA). Sections were cut from each tissue block at 4-μm thickness, placed on positively charged slides, and dried for 1 hour at 60°C ± 2°C. Excepting sample stability studies, all cut sections were tested within 2 months of sectioning.

#### Antibodies

All experiments were performed with monoclonal LAG-3 antibody 17B4 preparations manufactured from hybridoma cultures for Labcorp, except for analysis of clustered regularly interspaced short palindromic repeats (CRISPR)-engineered LAG-3 knockout cell lines, for which a commercially available LAG-3 17B4 antibody was obtained from LSBio (Cat. # LS-C18692) or as otherwise noted in the text.[22] For precision studies, 3 independent lots of antibody were produced from the 17B4 hybridoma. The working concentration of the LAG-3 17B4 antibody was 2.5 μg/mL. The negative control antibody, mouse monoclonal immunoglobulin G1 (IgG1) clone MOPC-21, was obtained from Leica Biosystems (Cat. # PA0996). Further details on the staining and melanin removal procedures are in the **supplemental material** and **supplemental table 1**.

### Melanin scoring

To determine the efficacy of the melanin removal step of the protocol, the amount of melanin pigment in the tumor region was scored on a scale of 0 to 4+. Definitions for melanin pigment scoring expected on melanoma tissue–stained slides and indications for the evaluability of the melanin interpretation in LAG-3 IHC assay scoring are provided in **supplemental table 2**.

### LAG-3 scoring

An overview of the LAG-3 scoring method is provided in **supplemental figure 1**. Evaluation criteria for staining intensity of LAG-3–positive ICs consisted of weak (1+), moderate (2+), and strong (3+) LAG-3–positive staining (**supplemental table 3**). In addition to cell-surface expression, LAG-3 protein is also retained in intracellular compartments.[23] Thus, LAG-3 IC positivity was quantified in cells that morphologically resembled lymphocytes with punctate (perinuclear and/or Golgi pattern), cytoplasmic, and/or membranous LAG-3 staining of any intensity above background (**supplemental figure 2**). LAG-3–positive IC content in the tumor region was visually estimated by microscopic examination by the study pathologists, following group alignment using a reference slide set. A hematoxylin and eosin-stained slide for each melanoma sample tested was reviewed by a pathologist to identify the overall tumor region and confirm the presence of ≥100 TCs. Results were reported as the percentage of LAG-3–positive ICs relative to all nucleated cells (ICs [lymphocytes and macrophages], stromal cells, and TCs) within the overall tumor region. The tumor region included TCs, intratumoral stroma, and peritumoral stroma (the band of stromal elements directly contiguous with the outer tumor margin). Normal and/or adjacent uninvolved tissues were not included (**supplemental figure 3**). The scoring scale was (in %) 0, 1, 2, 3, 4, 5, 10, and further increments of 10 up to 100. Samples with LAG-3–positive IC percentage scores of ≥1% were reported as LAG-3–positive.

The methods for the generation of CRISPR-engineered LAG-3 knockout cell lines, peptide inhibition assay, precision study measurements and reproducibility within the same laboratory and across laboratories, and stability experiments are provided in the **supplemental material**.

## RESULTS

### Components of the LAG-3 IHC assay

Primary antibody concentration and incubation times for assay components were optimized for appropriate positive staining, staining intensity, and overall staining quality of LAG-3 while minimizing nonspecific background staining. Antibody concentrations of 1.25 μg/mL, 2.5 μg/mL, 3.0 μg/mL, and 3.5 μg/mL were evaluated, and 2.5 μg/mL was determined to be the optimal concentration.

#### Detection of LAG-3 in tissues using the 17B4 clone antibody

To investigate the ability of the LAG-3 IHC assay to detect LAG-3 IC expression in human FFPE tissue samples, the assay was used to stain LAG-3 in commercially procured human tonsil tissue. We hypothesized that if the LAG-3 IHC assay detected LAG-3 IC expression, then staining would be present in lymphocytes, but not in nonimmune regions, such as the crypt epithelium. Staining of the tonsil tissues using the LAG-3 IHC assay revealed membranous/cytoplasmic staining of LAG-3 in lymphocytes in germinal center and interfollicular regions, but no LAG-3 staining in the crypt epithelium (**figure 1A**). Additionally, no staining was observed in the slide stained with the mouse IgG isotype control.

**FIGURE 1.**
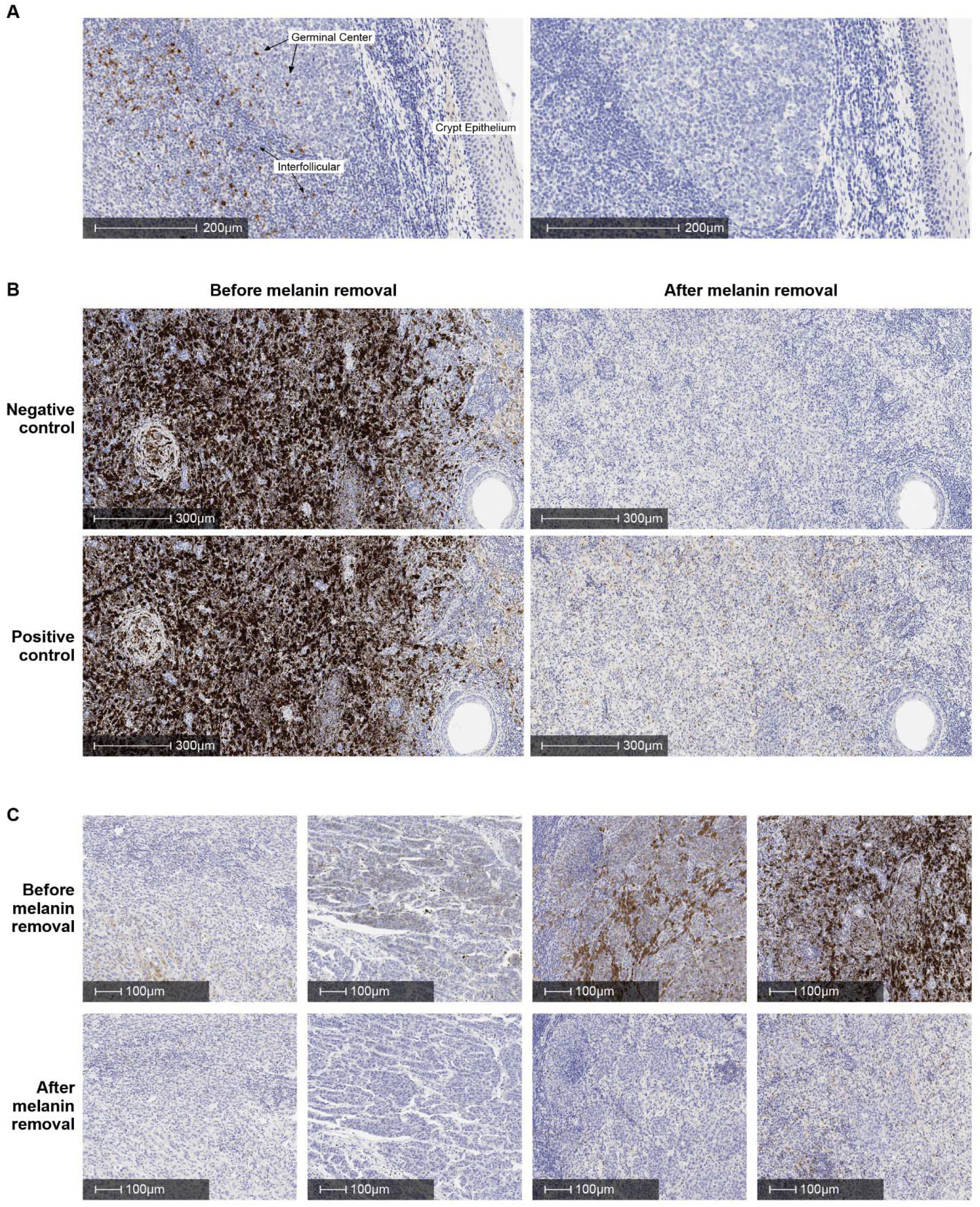
Identification of LAG-3 in human tissues using the LAG-3 IHC assay. A, Detection of LAG-3 in human tonsil tissue. Left-hand image depicts LAG-3 staining pattern in tonsil tissue showing moderate to strong plasma membrane/cytoplasmic staining in lymphocytes in germinal centers and interfollicular region. The crypt epithelium is negative. No staining is seen with negative reagent control (right-hand image). B, Staining of FFPE melanoma samples with negative reagent control (upper) or LAG-3 antibody (lower) before (left) and after (right) melanin removal procedure at 10× magnification. C, Examples of LAG-3 staining in FFPE melanoma samples before (upper) and after (lower) the melanin removal procedure at 20× magnification. FFPE, formalin-fixed paraffin-embedded; IHC, immunohistochemistry; LAG-3, lymphocyteactivation gene 3.

The LAG-3 IHC assay was developed to include attenuation of melanin staining from FFPE sections prior to IHC and to minimize the impact of melanin pigment on interpretation of the assay. Examples of different levels of melanin pigmentation are shown in **supplemental figure 4**. The efficacy of melanin removal from tissue samples using the melanin removal procedure is shown in **figures 1B** and **1C**. All melanoma tissue samples selected for further investigation had acceptable negative control staining and melanin pigmentation ≤1+. LAG-3 staining was consistent in bleached and unbleached serial sections from the same tissue block (data not shown).

### Specificity and sensitivity of the LAG-3 IHC assay

To investigate the specificity of the LAG-3 IHC assay, the *LAG3* gene was disrupted by CRISPR-mediated mutagenesis in COV434 cell lines. In total, 3 pooled cell lines were derived, each with differing levels of *LAG3* knockout (out-of-frame indel frequency = 71.02% in Cr1, 62.07% in Cr2, and 65.74% in Cr3) (**figure 2A**). The LAG-3 expression of these cell lines was compared with parental COV434 cells to investigate the specificity of the LAG-3 IHC assay. LAG-3 staining in parental COV434 cells was markedly higher than each of the 3 *LAG3* knockout cell lines, which each had staining consistent with anticipated levels of residual LAG-3 expression based on the frequency of alterations determined by next-generation sequencing (**figure 2B**). These data suggest that the LAG-3 IHC assay is specific for the detection of LAG-3 protein expression.

**FIGURE 2.**
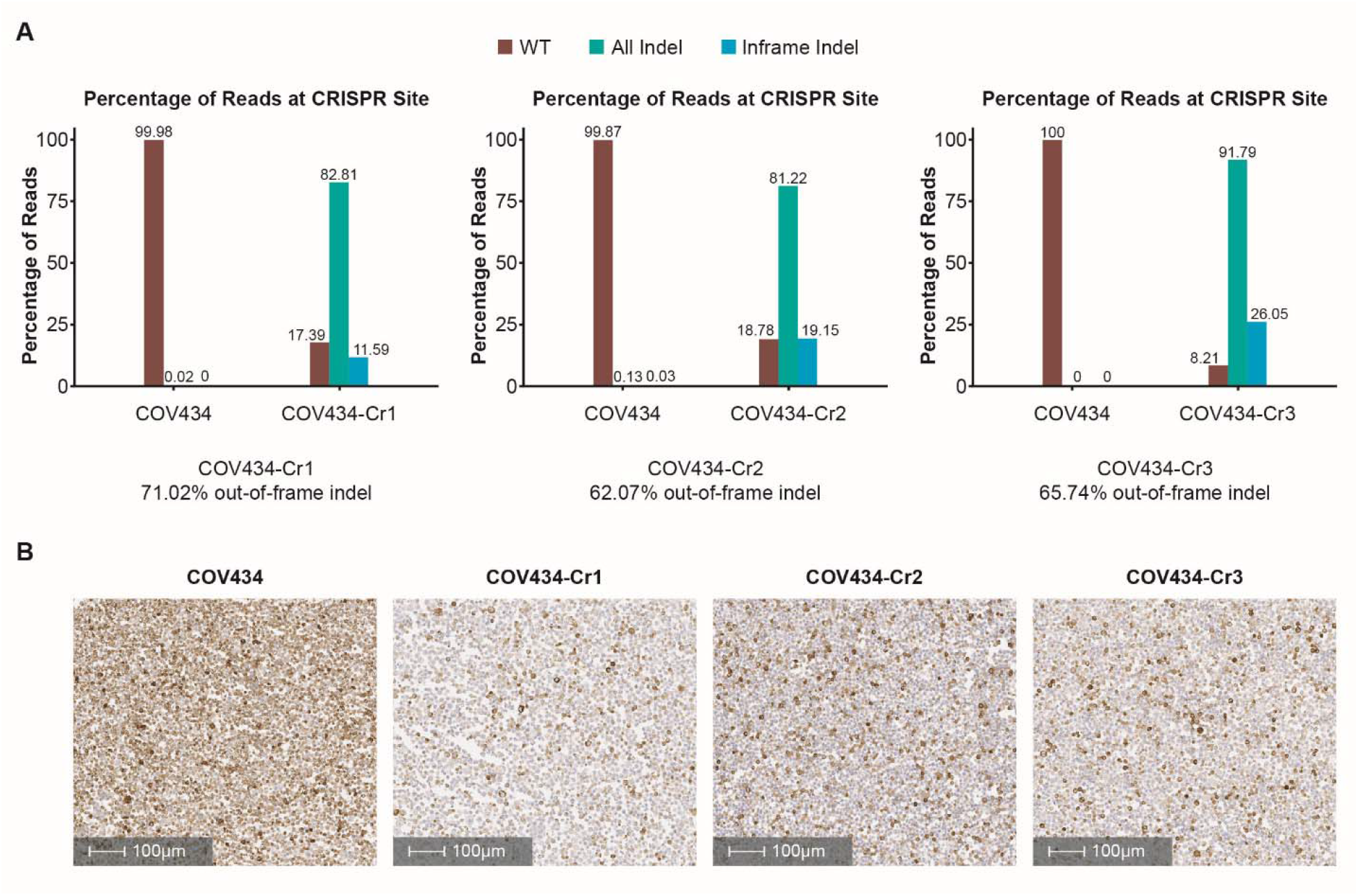
Detection of LAG-3 expression in parental COV434 cells and LAG-3–disrupted COV434 cells. A, Bar charts showing NGS results from each of the pooled CRISPR-engineered COV434 cell lines. B, IHC staining showing LAG-3 expression in parental COV434 cells and the 3 pooled CRISPR-engineered COV434 cell lines. Tonsil tissue was used as a positive/negative control for the IHC staining. CRISPR, clustered regularly interspaced short palindromic repeats; IHC, immunohistochemistry; LAG-3, lymphocyte-activation gene 3; NGS, next-generation sequencing; WT, wild type.

A peptide competition assay was performed using a synthetic LAG-3 peptide to further investigate the specificity of the LAG-3 IHC assay. The percentage of LAG-3–positive ICs in melanoma tissue was found to decrease from a starting staining level of 40% to <1% following preincubation with increasing molar ratios of a LAG-3 peptide (**table 1**), indicating that the LAG-3 peptide bound competitively to the 17B4 clone.

**TABLE 1.** LAG-3 IHC peptide competition validation results

| Specimen<br>(peptide:antibody ratio) | % LAG-3–positive<br>ICs | Staining intensity |
| --- | --- | --- |
| Melanoma LAG-3 mAb (0:1) | 40 | 2+ |
| Melanoma LAG-3 peptide (1:0) | 0 | N/A |
| Melanoma (1:1) | 40 | 2+ |
| Melanoma (2:1) | 30 | 2+ |
| Melanoma (5:1) | 10–20 | 1–2+ |
| Melanoma (10:1) | 2 | 1+ |
| Melanoma (30:1) | <1 | 1+ |

ICs, immune cells; IHC, immunohistochemistry; LAG-3, lymphocyte-activation gene 3; mAb, monoclonal antibody; N/A, not applicable.

To determine the range of LAG-3 IC expression in melanoma specimens, 100 commercially procured melanoma samples were assessed using the LAG-3 IHC assay. Of these 100 samples, 38 were positive for LAG-3 IC expression and 62 were negative, using 1% expression as a cutoff value (**figure 3**). The range of IC expression in the positive specimens was 1% to 40%, with a median of 3%. Of the positive cases, the majority (36) had a LAG-3 IC staining intensity of 2+, 1 sample had a LAG-3 IC staining intensity of 3+, and 1 sample had a LAG-3 IC staining intensity of 1+. Taken together, these data indicate that the LAG-3 IHC assay detects varying levels of immune infiltrates expressing LAG-3 in human FFPE melanoma samples. **Figure 4** shows representative tissue examples of staining from 0% to 30%.

**FIGURE 3.**
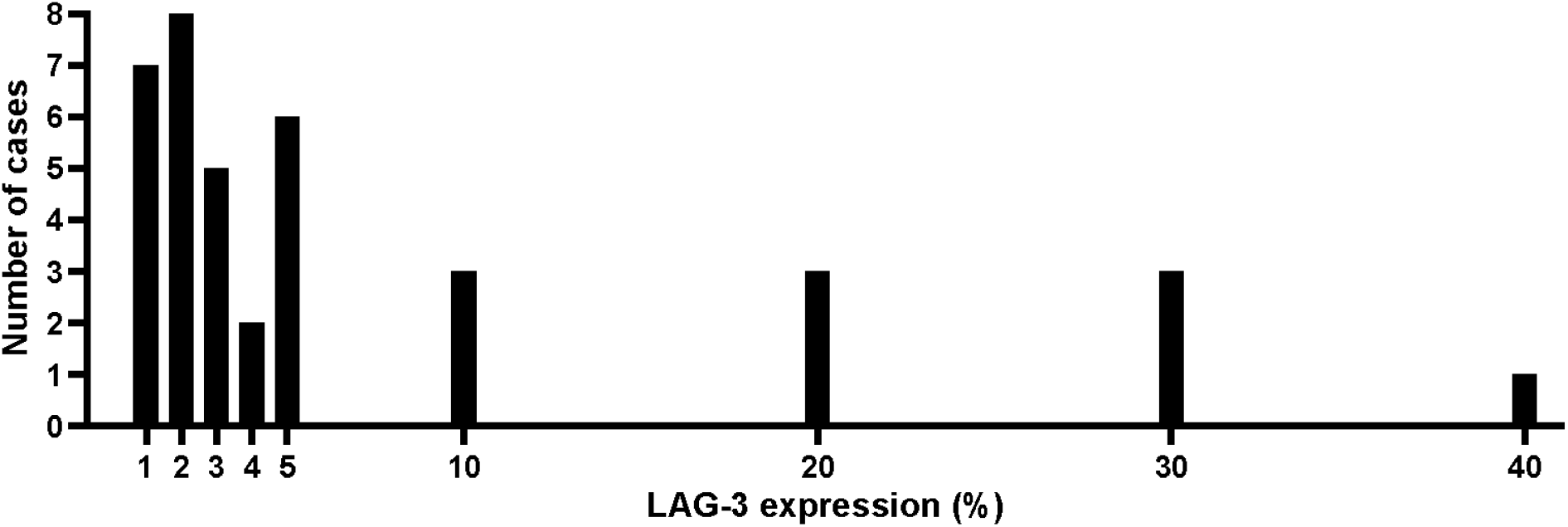
Detection of a range of LAG-3 expression levels using the LAG-3 IHC assay. Bar chart showing scoring distribution across LAG-3–positive samples (defined as those with LAG-3–positive IC content ≥1%) from a set of 100 commercially procured human FFPE melanoma specimens. Of the 100 samples, 38 were LAG-3–positive and 62 were LAG-3–negative. FFPE, formalin-fixed paraffin-embedded; IC, immune cell; IHC, immunohistochemistry; LAG-3, lymphocyte-activation gene 3.

**FIGURE 4.**
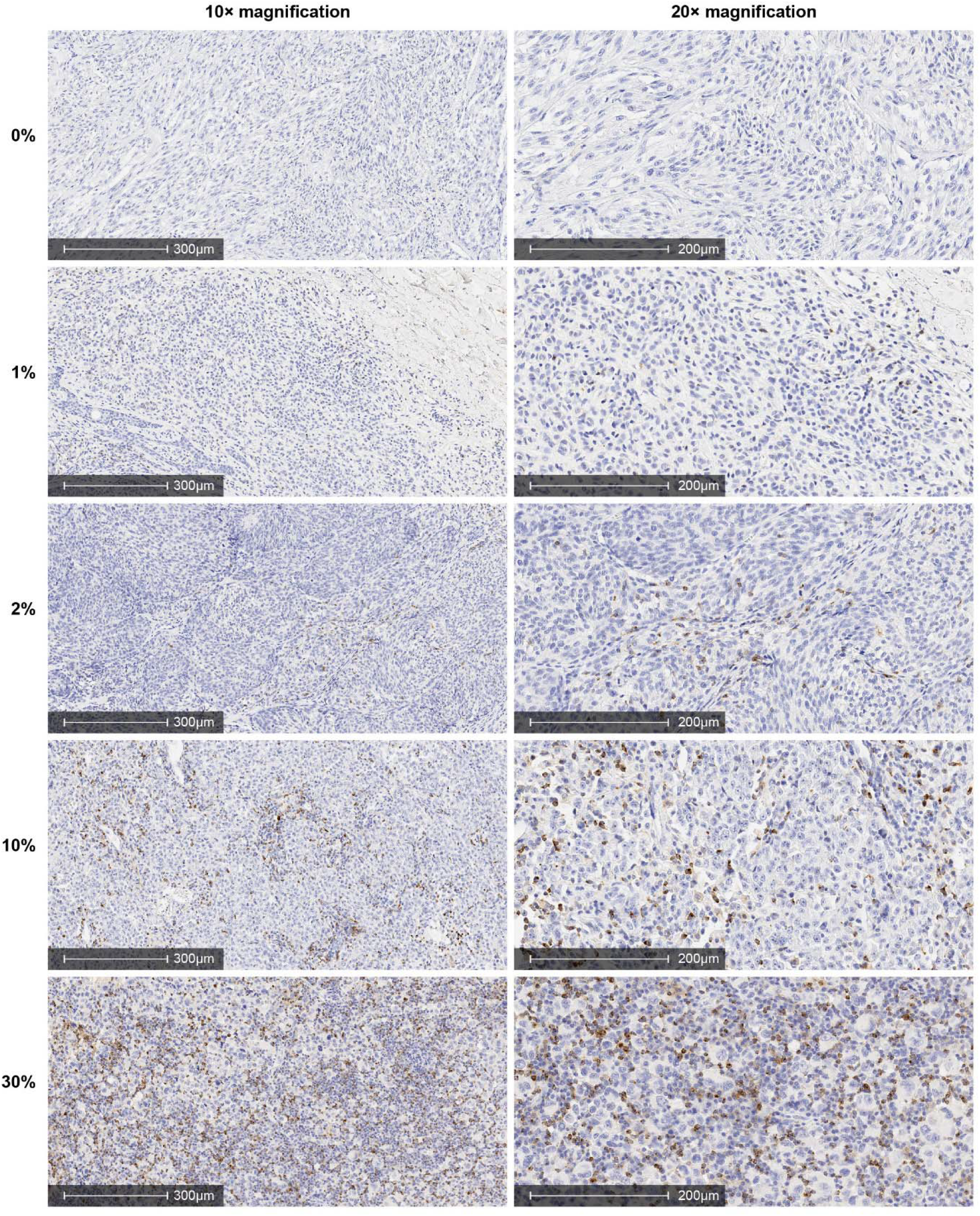
Examples of a range of LAG-3 expression levels detected in melanoma tissues using the LAG-3 IHC assay. Melanoma tissues showing a range of staining (0%–30%) for LAG-3 examined at magnifications of 10× (left-hand image) and 20× (right-hand image). IHC, immunohistochemistry; LAG-3, lymphocyte-activation gene 3.

### Analytical precision of the LAG-3 IHC assay within the same laboratory

Twenty-four FFPE melanoma samples and 1 normal human tonsil tissue control sample were stained on 2 different Leica BOND-III instruments and subsequently scored by 2 independent pathologists to establish the repeatability and reproducibility of the LAG-3 IHC assay. The intrarun repeatability, interday, interinstrument, interoperator, and interreagent lot reproducibility all demonstrated a high concordance, with all point estimates >95% in average negative agreement (ANA), average positive agreement (APA), and overall percentage agreement (OPA) (**table 2**).

**TABLE 2.** Summary of precision study results

| Evaluation | Percentage agreement (95% CI) |
| --- | --- |
|  | ANA: 98.5 (97.3–99.6) |
| Intrarun repeatability | APA: 98.6 (97.4–99.6) |
|  | OPA: 98.5 (97.3–99.6) |
|  | ANA: 97.4 (96.4–98.4) |
| Interday reproducibility | APA: 97.6 (96.6–98.5) |
|  | OPA: 97.5 (96.5–98.4) |
|  | ANA: 97.8 (96.8–98.6) |
| Interinstrument reproducibility | APA: 97.9 (97.0–98.7) |
|  | OPA: 97.8 (97.0–98.6) |
|  | ANA: 97.8 (96.8–98.6) |
| Interoperator reproducibility | APA: 97.9 (97.0–98.7) |
|  | OPA: 97.8 (96.9–98.7) |
|  | ANA: 97.4 (96.6–98.2) |
| Interreagent lot reproducibility | APA: 97.6 (96.8–98.3) |
|  | OPA: 97.5 (96.7–98.3) |

ANA, average negative agreement; APA, average positive agreement; CI, confidence interval; OPA, overall percentage agreement.

#### Interobserver and intraobserver reproducibility of the LAG-3 IHC assay within the same laboratory

Evaluations of 60 melanoma samples performed by 3 independent pathologists from the same laboratory and repeat evaluations of the same 60 melanoma samples by the same pathologist were examined to determine the interobserver and intraobserver reproducibility of the assay within the same laboratory. To determine the interobserver reproducibility of the LAG-3 IHC assay, pairwise comparisons were made of the 180 diagnostic calls by the 3 pathologists: 91 were concordant for positive-to-positive calls, and 77 were concordant for negative-to-negative calls. Disagreements occurred in 12 cases, all of which had LAG-3 scores around the 1% threshold (LAG-3–positive IC content of 0%–1%), resulting in a lower point estimate and lower bound 95% confidence interval (CI) for ANA compared with APA and OPA. Point estimates for ANA, APA, and OPA were >90% with the lower bound 95% CIs >85% (**table 3**).

**TABLE 3.**
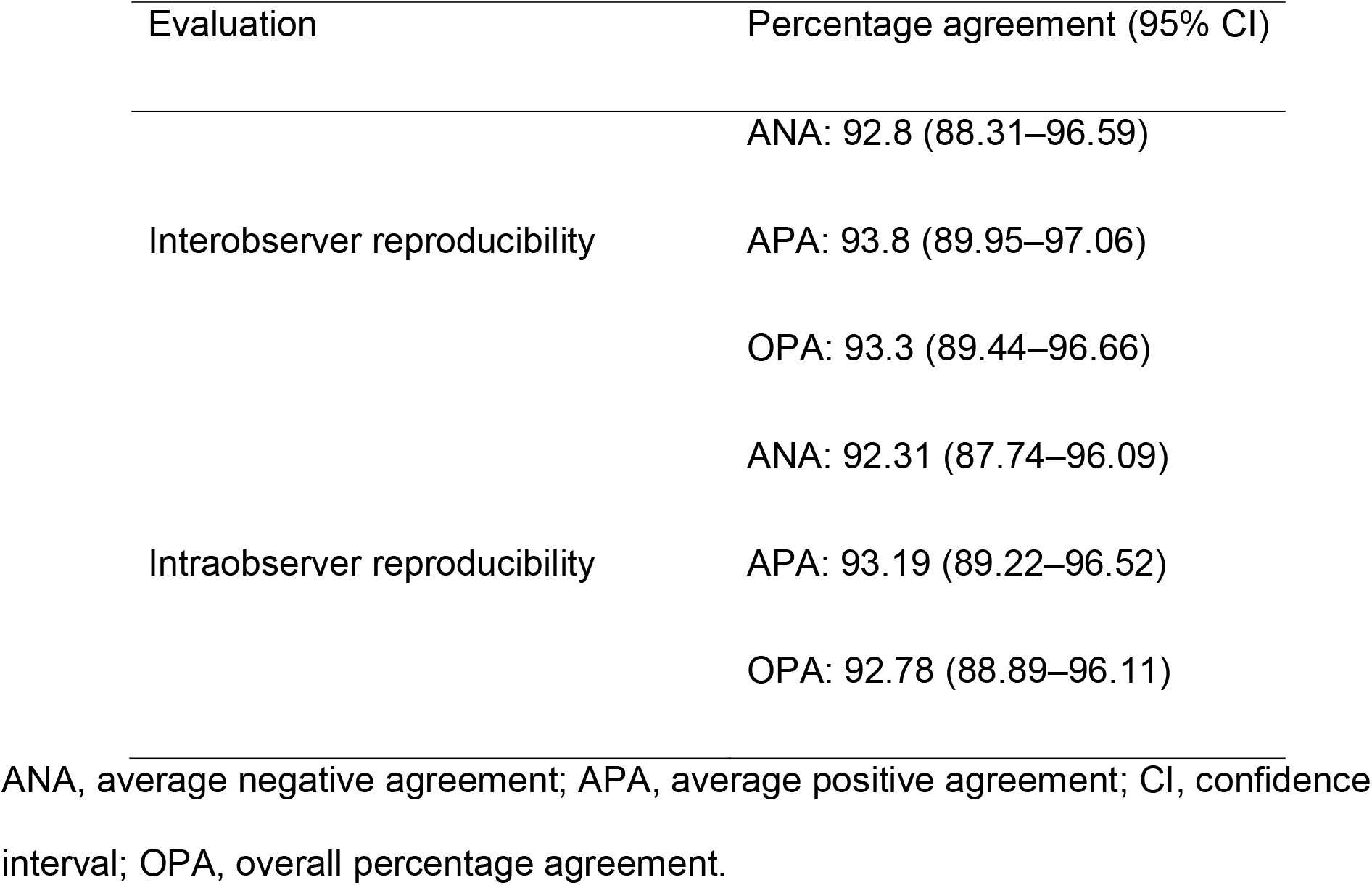
Percentage agreement and 95% CIs for interobserver and intraobserver agreement within the same laboratory

To determine intraobserver reproducibility of the LAG-3 IHC assay, the 60 samples assessed in the interobserver reproducibility testing were reassessed by the same pathologists, following a wash-out period. Among the 180 comparisons of diagnostic calls between 2 reads by 3 pathologists, 89 were positive-to-positive concordant, 78 were negative-to-negative concordant, 8 were negative-to-positive discordant, and 5 were positive-to-negative discordant. Additionally, the point estimates and lower bound 95% CIs were >90% and >85%, respectively, in ANA, APA, and OPA (**table 3**).

#### Interlaboratory and intralaboratory reproducibility of the LAG-3 IHC assay

Two experiments were performed to assess interlaboratory reproducibility: interobserver and intraobserver reproducibility, and overall interlaboratory and intralaboratory reproducibility. First, to investigate the interobserver and intraobserver reproducibility of the LAG-3 IHC assay between different laboratories, 70 melanoma LAG-3–prestained cases were assessed by 3 pathologists at 3 separate laboratories. Second, to determine overall interlaboratory and intralaboratory reproducibility, unstained slides from 24 melanoma cases that had previously been shown to have a range of LAG-3 expression were tested at 3 separate laboratories. The interobserver and intraobserver reproducibility and overall interlaboratory and intralaboratory reproducibility demonstrated assay staining and scoring concordance with point estimates for all studies at >90% in ANA, APA, and OPA and lower bound 95% CIs >85% (**table 4**).

**TABLE 4.**
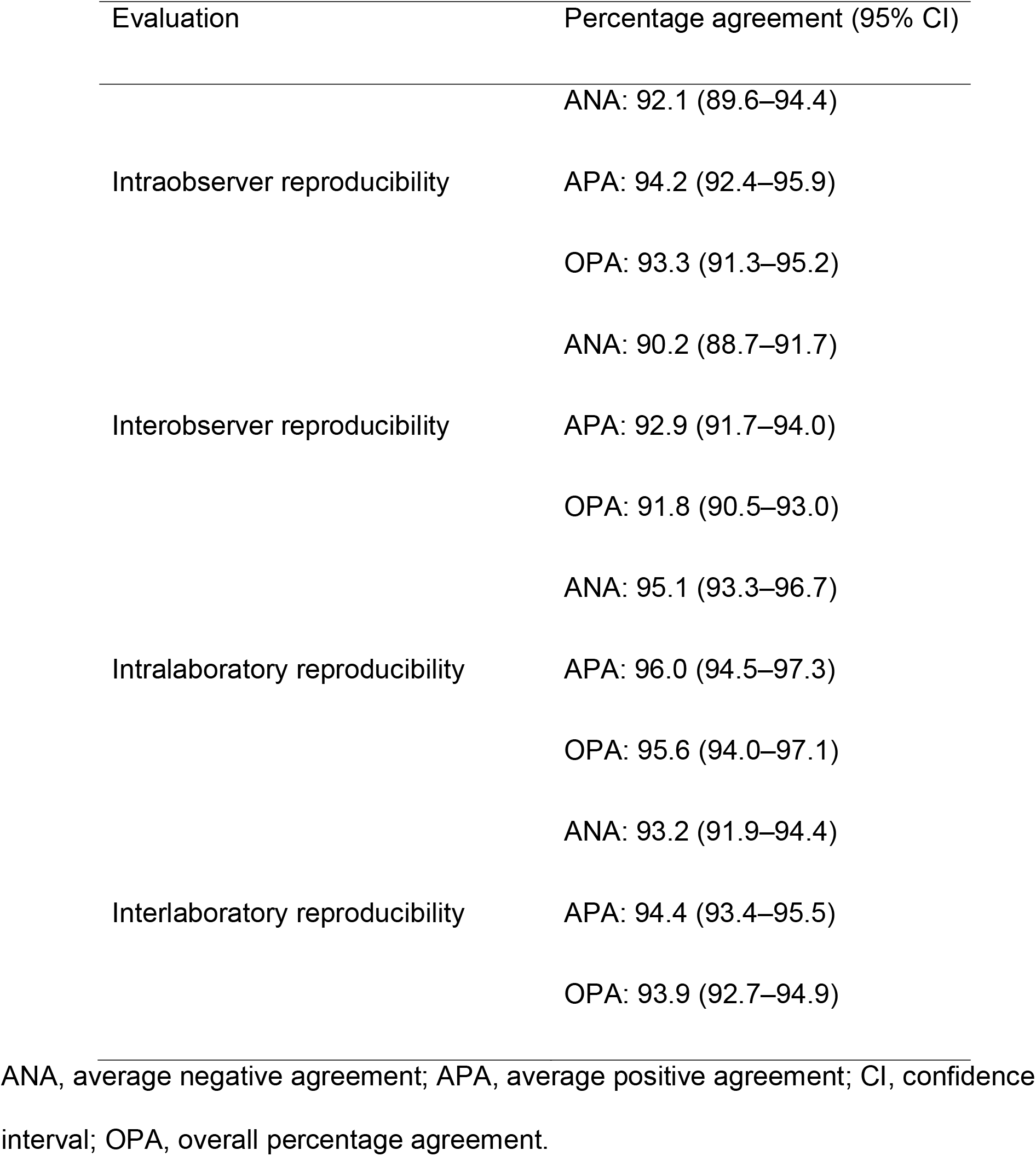
Percentage agreement and 95% CIs in the interlaboratory reproducibility study

### Slide stability experiments

To establish the stability of LAG-3 protein in unstained FFPE tissue sections on glass slides for the LAG-3 IHC assay, the concordance of sectioned tissue samples stained after different storage periods was measured. There was 100% concordance in scoring (positive or negative) at all time points for slides stored at ambient temperatures or 2–8°C. The LAG-3–positive IC staining intensity results for the tonsil tissue were 100% concordant from baseline through month 18 at both 2–8°C and ambient temperatures, with a decrease in LAG-3 IC staining intensity from 3+ to 2+ at month 24. Although there was some slight variation (increase or decrease) in the percentage of LAG-3–positive ICs for some melanoma samples during the course of testing (eg, a case reported as 2% at week 2, 1% at week 4, and 2% at month 2), the LAG-3 score (positive or negative) and LAG-3–positive IC staining intensity (1+, 2+, 3+) results were 100% concordant for individual samples tested at each time point and each temperature. The small differences observed may be attributable to variations in the density of ICs between tissue sections.

## DISCUSSION

LAG-3 is a key immune checkpoint currently being investigated as an I-O therapy for patients with solid tumors and hematological malignancies.[13, 16, 18, 21, 24–26] The development of a robust LAG-3 IHC assay will enable the analysis of IC LAG-3 status in the tumor microenvironment and the correlation between LAG-3 expression status and response to LAG-3–directed oncology treatments. A robust LAG-3 IHC assay that is suitable for clinical trials and clinical use for melanoma is described in this work. The specificity of the assay was demonstrated using cell lines with *LAG3* gene disruptions and with a peptide antigen competition assay. LAG-3 scoring was reported as the percentage of LAG-3–positive ICs (which morphologically resembled lymphocytes) relative to all nucleated cells within the overall tumor region. A ≥1% cutoff was used to determine LAG-3 positivity. Analytical precision was demonstrated for intrarun repeatability, interday, interinstrument, interoperator, and interreagent lot reproducibility, with concordance >95%. Pathologist interobserver and intraobserver reproducibility was >90% in terms of ANA, APA, and OPA. LAG-3 was observed to be stable in unstained tissues mounted on glass slides, with concordant staining observed in samples stored at both 2–8°C and ambient temperatures for up to 24 months. These data demonstrate that this assay can reproducibly determine the proportion of LAG-3–positive ICs within a sample. Despite challenges associated with the scoring of ICs, the LAG-3 IHC assay demonstrated a high level of interobserver reproducibility both within the same laboratory and between independent laboratories.[27, 28]

A particular issue for the interpretation of IHC assays for melanoma tissues is the presence of melanin pigment. Melanin pigmentation can interfere with IHC interpretation, as it may obscure morphological features and is similar in color to the chromogen 3,3’-diaminobenzidine tetrahydrochloride hydrate (DAB), which is commonly used in IHC assays, including the LAG-3 IHC assay described here. The pretreatment method described in this work removed melanin from samples without compromising the LAG-3 antigen and resulted in no samples that could not be interpreted due to excess melanin pigmentation.

One limitation of the studies presented in this work is that a number of preanalytical factors may impact the performance of the LAG-3 IHC assay, including location of the tissue assessed (ie, primary vs. metastatic),[29, 30] sample ischemia time, and fixation methods.[31] Additionally, the design of the cut slide stability studies compared LAG-3 staining and IC expression with baseline (time 0), but did not include comparison with other timepoints.

The assay described in this report was utilized to stratify patients based on LAG-3 expression in RELATIVITY-047 (NCT03470922), a phase 2/3 clinical trial in patients with previously untreated metastatic or unresectable melanoma. The trial compared combined nivolumab (anti–PD-1) and relatlimab (anti–LAG-3) treatment with nivolumab monotherapy, and benefit of combination therapy was observed in comparison with nivolumab monotherapy.[21] While the median PFS estimates were longer for patients with LAG-3 expression ≥1% across both treatment groups, a benefit with the combination therapy over nivolumab was observed regardless of LAG-3 expression. [21]

Both the present report and RELATIVITY-047 determined LAG-3 positivity using a ≥1% cutoff.[21] However, the prevalence of LAG-3 positivity observed in other sample sets or patient populations may vary, meaning cutoff values for clinical utility will have to be determined and validated in clinical studies. For instance, Dillon et al reported a higher prevalence of LAG-3 positivity using a ≥1% cutoff in a different set of commercially procured FFPE melanoma samples than in the melanoma samples used in this report.[32] Dillon et al also reported a higher prevalence of LAG-3 positivity in gastric and gastroesophageal cancer samples than in the melanoma samples used in this report. The LAG-3 assay described in this manuscript is currently being utilized in a number of clinical trials for multiple different tumor types.

In summary, a robust IHC assay for the determination of LAG-3 IC status in the tumor microenvironment in solid tumor tissues has been developed.

## Acknowledgments

The authors thank John Feder and Samantha Yost, both of Bristol Myers Squibb, for generating the CRISPR knock-out cell lines. Medical writing and editorial support were provided by Peter Harrison, PhD, and Matthew Weddig of Spark Medica Inc, funded by Bristol Myers Squibb.

## Authors’ Contributions

LJ, JY, BM, and JS designed the studies. LJ led the laboratory operation and procedures to provide stained slides to pathologists. BM was the lead pathologist for the study. BM, AS-C, SS, and KJ analyzed and interpreted the IHC slides and provided LAG-3 scores. JY provided statistical study design, data analyses, and interpretation. CS performed peptide inhibition assay. SA reviewed the data and provided input on the interpretation of the data. JS, CS, LJ, LD, CH, and JW provided input on data analysis and interpretation. LD co-led LAG-3 IHC diagnostic development with Labcorp. LD, JW, and CH developed the validation strategy, in partnership with Labcorp, and reviewed and approved the experimental design and validation reports. JW and CH served as pathology subject matter experts for LAG-3 IHC assay development. DL oversaw assay verification and optimization experiments in support of assay transfer to Labcorp and trained Labcorp staff on using the LAG-3 IHC assay. CH trained pathologists at Labcorp on manual scoring of the LAG-3 IHC assay and developed the LAG-3 IHC scoring algorithm and the assay scoring manual used at Labcorp. All authors contributed to drafting, reviewed, and approved the manuscript.

## Data availability statement

The datasets generated during and/or analyzed during the current study are not publicly available but are available from the corresponding author on reasonable request.

