## Supplemental Material for "Development of a LAG-3 Immunohistochemistry Assay for Melanoma"

### SUPPLEMENTAL METHODS

#### Staining procedures

Formalin-fixed, paraffin-embedded (FFPE) tissue sections were deparaffinized by immersing slides through xylene and graded alcohols prior to staining. Variable amounts of melanin pigment are typically found in melanoma tumors, potentially impacting the interpretation of the lymphocyte-activation gene 3 (LAG-3) immunohistochemistry (IHC) assay. To attenuate melanin pigment in the melanoma samples, deparaffinized slides were placed in melanin-removal agent (1 part Dako Target Retrieval solution pH 9, 10× concentrate [Agilent, Cat. # S236784-2], 8 parts methanol with 1 part hydrogen peroxide, 30% w/w, added) and incubated in a Decloaking Chamber™ NxGen (BioCare Medical, Pacheco, CA; part no. DC2012) for 3 hours at 60°C, then rinsed in deionized water.

Following melanin removal (“bleaching”), the LAG-3 IHC assay was performed in the BOND-III automated staining system (Leica Biosystems) as follows (summarized in **supplemental Table 1**):

1. Antigen unmasking of the FFPE tissue sections was done by incubating samples in BOND™ Epitope Retrieval Solution 1 (Leica Biosystems, Cat. # AR9961) for 20 minutes at 100°C

2. Endogenous peroxidase activity was quenched by incubation in pre-primary peroxidase inhibitor (BOND™ Polymer Refine Detection; Leica Biosystems, Cat. # DS9800) for 5 minutes at ambient temperature (20–25°C)
3. Slides were incubated with a protein block (Dako Serum Free Protein Block; Agilent, Cat. # X090930-2) for 5 minutes and then incubated with the LAG-3 primary antibody, diluted to 2.5 µg/mL (1:400) in BOND Primary Antibody Diluent (Leica Biosystems, Cat. # AR9352) or negative control antibody (mouse monoclonal immunoglobulin G1 [IgG1]; clone MOPC-21; Leica Biosystems, Cat. # PA0996) for 30 minutes at ambient temperature
4. The primary antibody was washed off and the slides incubated with the post-primary rabbit anti-mouse immunoglobulin G (IgG) linker reagent (BOND™ Polymer Refine Detection; Leica Biosystems, Cat. # DS9800) for 8 minutes at ambient temperature
5. Incubation with the secondary polymer anti-rabbit poly–horseradish peroxidase-IgG was done for 8 minutes, followed by incubation with the 3,3'-diaminobenzidine tetrahydrochloride hydrate (DAB) chromogen (BOND™ Polymer Refine Detection; Leica Biosystems, Cat. # DS9800) for 10 minutes at ambient temperature
6. After washing off the excess DAB, sample nuclei were counterstained with hematoxylin (BOND™ Polymer Refine Detection; Leica Biosystems, Cat. # DS9800) for 5 minutes at ambient temperature

Melanoma tissue staining was performed with 3 run controls: melanoma biopsy tissue with LAG-3 immune cell (IC) expression >5% (predetermined by IHC) was used as a positive control, tonsil tissue with areas differentiated by positive or negative LAG-3 IC expression was used as a positive and negative control, and nonimmune mouse IgG was used as a negative reagent control. Slides were reviewed by a pathologist using bright field microscopy. If either the positive melanoma tissue control or the tonsil tissue control was deemed unacceptable by the interpreting pathologist, the staining run was repeated. To be considered acceptable, the positive melanoma tissue control must have had a LAG-3 IC expression score (see **LAG-3 scoring** section) of >5%, and the tonsil tissue must have had positive staining on ICs in germinal centers or interfollicular regions with no staining within the crypt epithelium, skeletal and smooth muscle fibers, collagen fibers, adipose tissue, and peripheral nerves.

**Generation of CRISPR-engineered LAG-3 knockout cell lines**

Pooled clustered regularly interspaced short palindromic repeats (CRISPR)-engineered COV434 ovarian granulosa tumor cells (TCs) with heterogeneous LAG-3 expression were derived using 3 unique, nonoverlapping CRISPR guides targeting different regions of exon 2 of LAG-3. Guide sequences were: Cr1, TGACCCCTGCTCTTCGCAGA; Cr2, GATCCTGGAGGGGGATTGTG; Cr3, GCCAGGGGCTGAGGTCCCGG. Editing frequency was assessed by next-generation sequencing. In total, 3 cell line pools were derived with different frequencies of modification, which led to absence of LAG-3 protein expression but not LAG-3 mRNA expression.

**Peptide inhibition assay**

A peptide corresponding to the immunogen used to generate antibody 17B4, GPPAAAPGHPLAPGPHPAAPSSWGPRPRRY, was synthesized and combined in various molar ratios (0-fold, 1-fold, 2-fold, 5-fold, 10-fold, and 30-fold excess) with antibody 17B4 in phosphate buffer saline solution for 30 minutes at ambient temperature and centrifuged at 16,000 × g at 4°C for 15 minutes. Supernatants from centrifuged aliquots were used as the primary antibody solution in the LAG-3 IHC assay performed on FFPE melanoma tissue previously scored with >5% LAG-3–positive ICs.

**Precision Study Measurements**

The agreements of LAG-3 scores were assessed to determine the intrarun repeatability and interday, interinstrument, interoperator, and interreagent lot reproducibility. Samples for this study consisted of 1 normal human tonsil to serve as both a positive and negative control and 24 FFPE melanoma tissues previously confirmed to have a range of LAG-3 IC expression (12 were LAG-3–positive [≥1%] with a range of 1%–40%, and 12 were LAG-3–negative [<1%]). Slides were sectioned from each FFPE melanoma tissue block as described above. One slide from each of the 24 melanoma tissue blocks was stained for hematoxylin and eosin (H&E). Each sample was tested on 5 nonconsecutive days, with 2 independent runs by 2 different operators on each day following a wash-out period.

**Intrarun Repeatability**

Intrarun duplicates were included in 2 independent runs per sample each day. All slides were evaluated by 2 pathologists. Each pathologist had 240 intrarun duplicates (24 specimens, ran twice each day for 5 days) evaluated for agreement, and 2 pathologists had a total of 480 combined pairwise comparisons to compute average negative agreement (ANA), average positive agreement (APA), and overall percentage agreement (OPA).

**Interday Reproducibility**

Two independent runs were performed each day. Each sample was run in duplicate and evaluated by 2 different pathologists who consolidated their evaluations into 1 call representing the run (agreed or discordant). For each pathologist or run, there were 10 interday pairwise comparisons per specimen (5 × 2). For both pathologists and runs, there were 20 interday pairwise comparisons (5 × 2 × 2). There were 480 (24 × 10 × 2) interday pairwise comparisons from 1 pathologist and 960 interday pairwise comparisons from 2 pathologists' reads to be evaluated for ANA, APA, and OPA.

**Interinstrument Reproducibility**

Two different Leica BOND-III instruments were used by each operator each day during the interday reproducibility testing; due to the limited number of slides allowed on each instrument, 12 specimens were run on 1 instrument while the other 12 were run on the other instrument. Because of the design and rotation of 2 instruments in 5 testing days, the number of runs on 2 instruments for 1 specimen were 6 and 4. The number of total interinstrument pairwise comparisons were 24 per specimen (6 × 4), 576 for all 24 specimens (24 × 24) for 1 pathologist, and 1152 (2 × 576) to be evaluated in total for 2 pathologists' reads. The total 1152 pairwise interinstrument comparisons were used to compute ANA, APA, and OPA.

**Interoperator Reproducibility**

Each sample was evaluated in 2 independent runs by 2 different operators each day. The number of runs by each operator was 5 for each sample. The total number of pairwise comparisons between the 2 operators per sample was 25 (5 × 5) and 600 (24 × 25) pairwise comparisons for all 24 specimens for each pathologist's read. In total, there were 1200 interoperator pairwise comparisons for 2 pathologists. The 1200 pairwise comparisons were used to compute ANA, APA, and OPA.

#### Interreagent Lot Reproducibility

Three reagent lots were used in rotation of 2 lots for each of the 5 testing days during the interday reproducibility testing, resulting in 4 runs with the first 2 lots and 2 runs with the third lot. The interreagent lot pairwise comparisons were 32 per specimen ([4 × 4] + [4 × 2] + [4 × 2]), adding up to 768 for all 24 specimens (32 × 24) for each pathologist's evaluation. In total, 1536 (2 × 768) interlot pairwise comparisons were used to compute ANA, APA, and OPA.

### Reproducibility Within the Same Laboratory

#### Interobserver Reproducibility

Sixty melanoma samples with a range of staining intensity and a minimum of 15% of challenging cases around the prespecified threshold (≥1%) were assessed by 3 independent, board-certified anatomic pathologists from the same laboratory. Samples were randomized and blinded prior to evaluation. Three pairwise comparisons were made and pooled to estimate ANA, APA, and OPA.

#### Intraobserver Reproducibility

The same 60 samples used to determine interobserver reproducibility were re-evaluated by the same 3 pathologists, following a wash-out period between original evaluations and re-evaluations. Results from the re-evaluations were compared with the original evaluations to assess intraobserver precision. Pathologists were blinded to original results, and slides were re-randomized prior to examination. Three intraobserver pairwise comparisons were made for each of the 3 pathologists and pooled to provide average intraobserver agreement from all 3 pathologists.

### Reproducibility Across Independent Laboratories

#### Interobserver and Intraobserver Reproducibility in Different Laboratories

Seventy melanoma samples with a range of staining intensity were assessed by 3 pathologists from 3 separate laboratories. Assessment occurred over 3 days at least 14 days apart, with 210 reads per pathologist. For intraobserver reproducibility, ANA, APA, and OPA were computed using all non-redundant pairwise comparisons for a single observer. For interobserver reproducibility, all non-redundant pairwise comparisons between pathologists (including laboratory 1 vs. laboratory 2, laboratory 1 vs. laboratory 3, and laboratory 2 vs. laboratory 3) were used to compute ANA, APA, and OPA.

#### Interlaboratory and Intralaboratory Reproducibility

Twenty-four melanoma cases with a range of LAG-3 IHC expression were tested on 5 different days at each of the 3 different laboratories. Intralaboratory ANA, APA, and OPA were computed using a pool of all possible nonredundant pairwise intralaboratory comparisons. Interlaboratory ANA, APA, and OPA were calculated using a pool of all possible nonredundant pairwise interlaboratory comparisons.

*Statistical Methods*

ANA, APA, and OPA were calculated for intrarun repeatability and interday, interinstrument, interreagent lot, interobserver and intraobserver, and interlaboratory and intralaboratory reproducibility measurements. 95% confidence intervals were calculated using the percentile bootstrap method.[1]

**Stability Experiments in FFPE Sections**

Slides were sectioned from 6 melanoma FFPE tissue blocks spanning the dynamic range of LAG-3 expression and 1 tonsil FFPE tissue block as described in the ***Tissue Specimens*** section. Half of the slides were stored at ambient temperature, and half were stored at 2–8°C. Two of the slides stored at ambient temperature, and 2 of the slides stored at 2–8°C were used for testing at different time periods: at time 0 (baseline), at 1, 2, and 4 weeks, and then at 2, 3, 4, 5, 6, 8, 10, 12, 14, 18, and 24 months. Using the LAG-3 IHC assay, 1 slide was stained with LAG-3 antibody and 1 slide with nonimmune mouse IgG. A tonsil tissue control was included with each staining run as a positive and negative control, as described in the **Staining Procedures** section.

**SUPPLEMENTAL TABLE 1.** Summary of the LAG-3 IHC Assay Staining Procedure Steps.

| Step | Procedure | Process | Reagent |
|---|---|---|---|
| 1 | Antigen retrieval | BOND™ Epitope Retrieval Solution 1, 20 min, 100°C | Leica Biosystems, Cat. # AR9961 |
| 2 | Pre-primary peroxidase activity inhibition | BOND™ Polymer Refine Detection,<br>5 min, ambient temperature | Leica Biosystems, Cat. # DS9800 |
| 3 | Protein block | Dako Serum Free Protein Block, 5 min | Agilent, Cat. # X090930-2 |
| | Primary antibody | Clone 17B4 (2.5 µg/mL) in BOND™ Primary Antibody Diluent or negative control antibody, 30 min, ambient temperature | Labcorp (antibody) Leica Biosystems (antibody diluent), Cat. # AR9352 |
| 4* | Post-primary rabbit anti-mouse IgG linker | BOND™ Polymer Refine Detection,<br><10 µg/mL in 10% (v/v) animal serum in TBS/0.1% ProClin™ 950, 8 min | Leica Biosystems, Cat. # DS9800 |

| | | | |
|---|---|---|---|
| 5 | Polymer anti-rabbit poly-HRP-IgG<br><br>DAB chromogen | BOND™ Polymer Refine Detection, 8 min<br><br>BOND™ Polymer Refine Detection, 66 mM in stabilizer solution, 10 min | Leica Biosystems, Cat. # DS9800<br><br>Leica Biosystems, Cat. # DS9800 |
| 6† | Hematoxylin counterstain | BOND™ Polymer Refine Detection, 5 min | Leica Biosystems, Cat. # DS9800 |

*Remove primary antibody by washing prior to this step.

†Remove excess DAB by washing prior to this step.

DAB, 3,3’-diaminobenzidine tetrahydrochloride hydrate; HRP, horse-radish peroxidase; IgG, immunoglobulin G; min, minutes; IHC, immunohistochemistry; LAG-3, lymphocyte-activation gene 3; TBS; tris-buffered saline; v/v, volume/volume.

**SUPPLEMENTAL TABLE 2.** Melanin Interpretation and Scoring Criteria.

| Interpretation | Staining Description | LAG-3 IHC Assay Scoring Acceptability |
|---|---|---|
| 0 | No melanin pigment observed | Yes |
| 1+ | 1 to 2 small foci in melanin containing tumor cells or macrophages | Yes |
| 2+ | More than 2 small foci of moderate to strong melanin or diffuse weak melanin with sufficient areas not obscured by melanin | Yes |
| 3+ | Diffuse weak to moderate melanin obscuring a significant portion of the tumor region | No |
| 4+ | Diffuse moderate to strong melanin obscuring most of the tumor region | No |

IHC, immunohistochemistry; LAG-3, lymphocyte-activation gene 3.

**SUPPLEMENTAL TABLE 3.** LAG-3 Overall Stain Intensity Interpretation Criteria.

| Interpretation | Staining Description |
|---|---|
| 1+ | Weak LAG-3–positive IC staining: light brown or very punctate staining that may require high-power (40×) examination to detect |
| 2+ | Moderate LAG-3–positive IC staining: moderate to dark brown staining that is easily visible with 20× objective |
| 3+ | Strong LAG-3–positive IC staining: dark brown staining that is easily visible with 10× or 20× objective and obscures cell detail |

IC, immune cells; LAG-3, lymphocyte-activation gene 3.

**SUPPLEMENTAL FIGURE 1.** LAG-3 IC scoring method overview. H&E, hematoxylin and eosin; IC, immune cells; LAG-3, lymphocyte-activation gene 3.

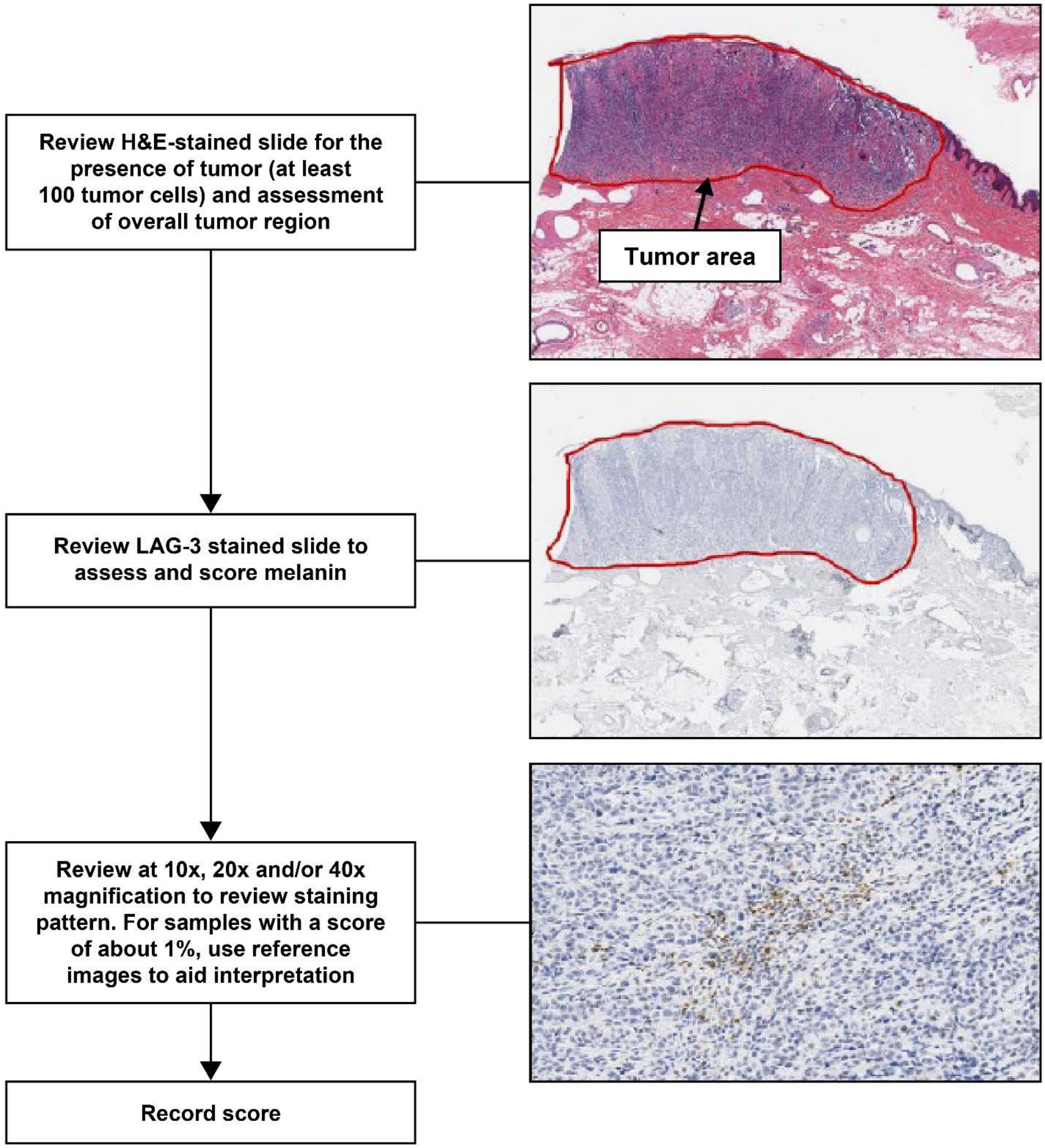

**SUPPLEMENTAL FIGURE 2.** Examples of punctate, membrane, and cytoplasmic LAG-3 IC staining observed with the LAG-3 IHC assay. Image shown at 40× magnification. IC, immune cells; IHC, immunohistochemistry; LAG-3, lymphocyte-activation gene 3.

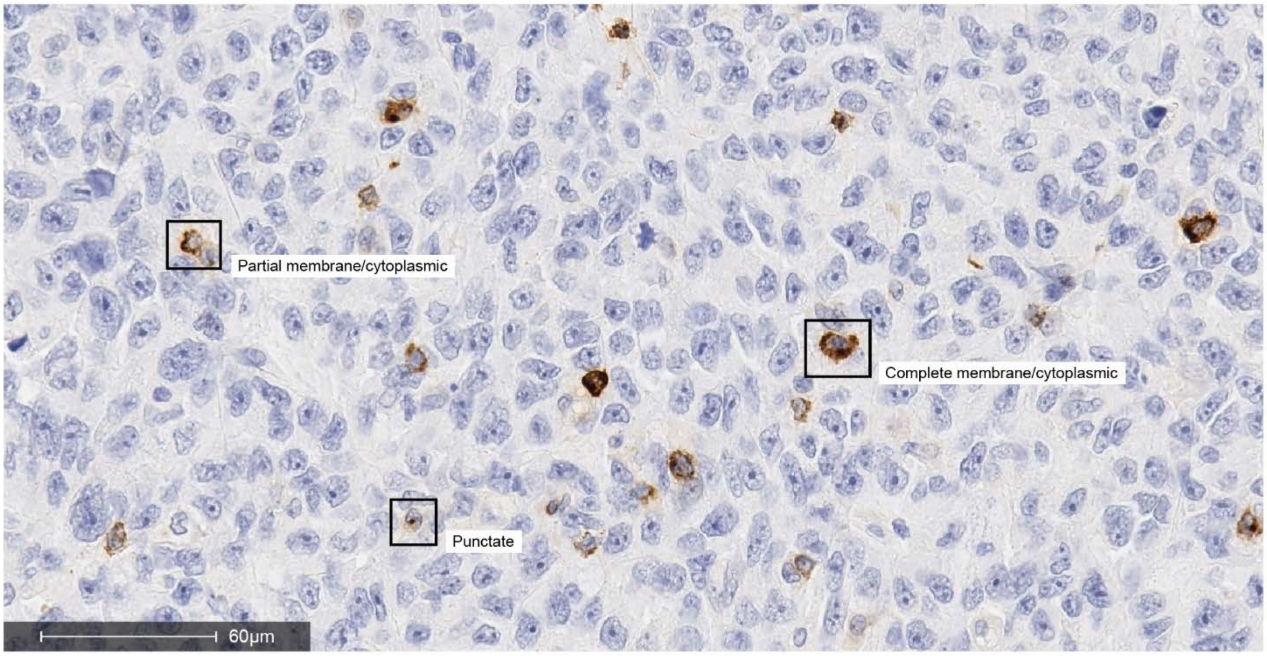

**SUPPLEMENTAL FIGURE 3.** Examples of the regions of a slide that were scored or not scored. A, Melanoma tissue with LAG-3–stained lymphocytes. The area scored includes the TC area and the PTS (outlined in red on the left panel). Adjacent normal (N) or uninvolved areas (shaded in pink) were not scored. Left panel image is shown at 10× magnification, right panel image is shown at 40× magnification. B, H&E–stained melanoma metastatic in lymph node. The area scored is shaded in blue (left panel) and included the TC area and PTS. Adjacent LN, shaded in pink (left panel), was not scored. The image is shown at 10× magnification. H&E, hematoxylin and eosin; LAG-3, lymphocyte-activation gene 3; LN, lymph node; PTS, peritumoral stroma; TC, tumor cell.

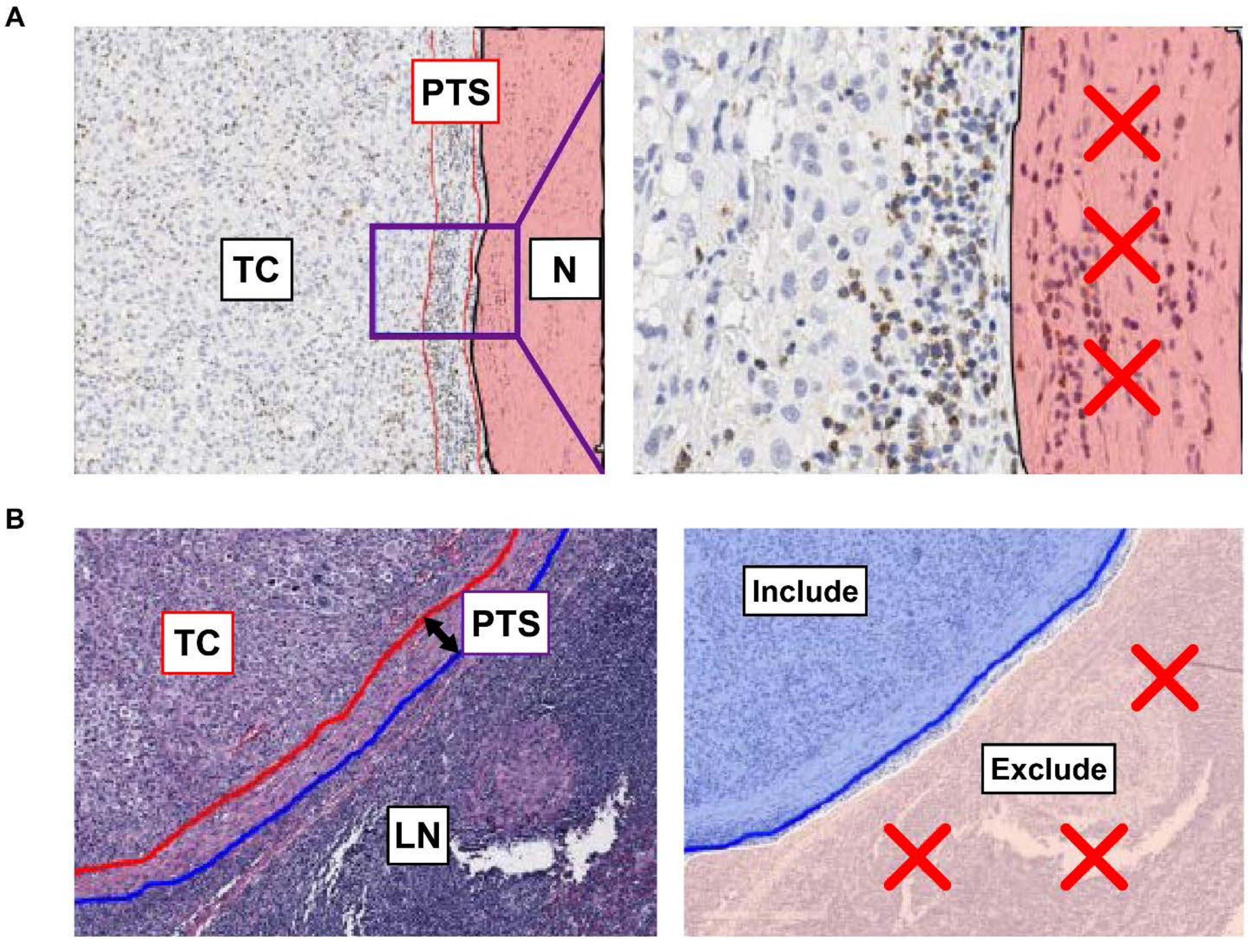

**SUPPLEMENTAL FIGURE 4.** Examples of LAG-3 IC staining in melanoma samples containing various levels of melanin pigment. A, Example of 1+ melanin pigmentation showing a single focus of tumor cells and macrophages with melanin. Image shown at 1× magnification (inset at 20×). B, Example of 2+ melanin pigmentation showing 2 large areas containing pigmented tumor cells and macrophages (outlined in red) showing weak to moderate melanin. Left-hand image shown at 2× magnification and right-hand image at 10×. C, Example of 3+ melanin pigmentation showing diffuse areas of moderately pigmented tumor cells and macrophages (outlined in red). An area with only limited melanin is present (outlined in green). Left-hand image shown at 0.7× magnification and right-hand image at 10×. D, Example of 4+ melanin pigmentation showing diffuse areas of strongly pigmented tumor cells and macrophages. LAG-3–stained lymphocytes cannot be visualized in the entire sample. Left-hand image is shown at 2× magnification and right-hand image at 10×. IC, immune cells; LAG-3, lymphocyte-activation gene 3.

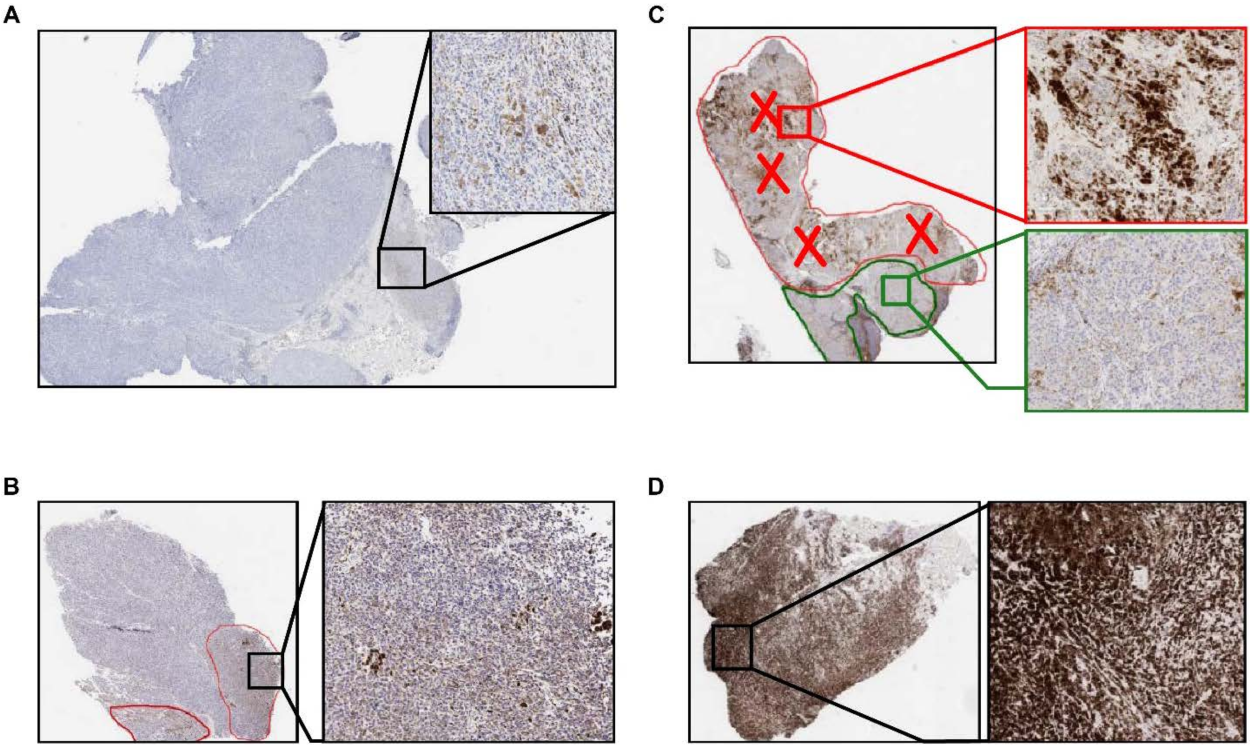

A
B
C
D
